# Tuning the Pennes Perfusion Rate to Model Large Vessel Cooling Effects in Hepatic Radiofrequency Ablation

**DOI:** 10.1101/2021.10.19.464935

**Authors:** Nikhil Vaidya, Marco Baragona, Valentina Lavezzo, Ralph Maessen, Karen Veroy

## Abstract

Radio-frequency ablation (RFA) has become a popular method for the minimally invasive treatment of liver cancer. However, the success rate of these treatments depends heavily on the amount of experience the clinician possesses. Mathematical modelling can help mitigate this problem by providing an indication of the treatment outcome. Thermal lesions in RFA are affected by the cooling effect of both fine-scale and large-scale blood vessels. The exact model for large-scale blood vessels is advection-diffusion, i.e. a model capable of producing directional effects, which are known to occur in certain cases. In previous research, in situations where directional effects do not occur, the advection term in the blood vessel model has been typically replaced with the Pennes perfusion term, albeit with a higher-than usual perfusion rate. Whether these values of the perfusion rate appearing in literature are optimal for the particular vessel radii in question, has not been investigated so far. The present work aims to address this issue. An attempt has been made to determine, for values of vessel radius between 0.55 mm and 5 mm, best estimates for the perfusion rate which minimize the error in thermal lesion volumes between the perfusion-based model and the advection-based model. The results for the best estimate of the perfusion rate presented may be used in existing methods for fast estimation of RFA outcomes. Furthermore, the possible improvements to the presented methodology have been highlighted.

## 1 Introduction

Liver cancer is among the most common cancers in the world with a high overall mortality-to-incidence ratio of 0.6 [1]. For its treatment, minimally invasive methods are preferred to surgical resection which is not suitable for patients with certain co-morbidities [2]. Among minimally invasive methods of treatment, such as cryoablation, high-intensity focussed ultrasound, radio-frequency ablation (RFA) etc. RFA is the most widely used [3]. During RFA an electrode is inserted into the tumour which sets up a radio-frequency current and causes thermal necrosis of the neighbouring tissue [2]. The challenge is that success rate still depends on the amount of experience the clinician possesses [4]. The liver is a highly perfused organ and RFA thermal lesions are affected by the cooling effect of fine-scale vessels (diameter smaller than 1 mm) as well as large vessels. The distinction between small and large vessels is based on the typical resolution of 1 mm of current CT scans. Previous research by Shrivastava et al. has indicated that the thermal significance of blood vessels in RFA depends on the temperature at the blood inlet and the surrounding tissue temperature, but not on their diameters [5].

Mathematical modelling can help solve this problem by providing an indication of treatment outcome. A number of mathematical models have been proposed for this purpose, which make a distinction between fine-scale vessels and large vessels [6, 7]. Fine vessels are not resolved in the simulation geometry and a volumetric term (e.g. Pennes perfusion term [8]) in the tissue subdomain is used to model their cooling effect. Large vessels, on the other hand, have strong localized cooling effects and need to be resolved fully in the simulation geometry. Additional terms modelling their cooling effects, appear only in the large vessel subdomains. Large vessels have been found to have pure cooling effects or directional effects depending on their size [9, 10].

The exact mathematical model for large blood vessels is advection-diffusion [6, 7]. The advection term present is capable of producing directional effects (if they occur for the particular blood vessels in question). However, in finite element simulations, the advection term is difficult to deal with and requires additional stabilization. We have reported in a previous publication that for a vessel radius larger than 0.5 mm, assuming the flow-rate to be given by Murray’s Law, no directional stretching of the thermal lesion in the flow direction should be expected, indicating a pure cooling effect [9]. In the past, the pure cooling effect of blood vessels has been modelled by replacing the advection term with the Pennes perfusion term *ωρ_b_C_b_(T − T_0_)* (albeit with a higher than usual perfusion rate *ω*) inside the blood-vessel region [11, 12]. This term is symmetric in the weak form of the PDE and also does not require additional stabilization. The following values of *ω* were used by Kröger et al. [11]:

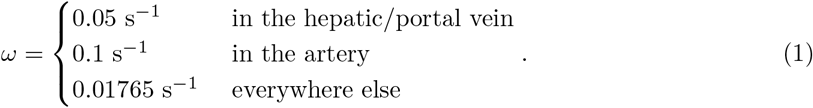

Audigier used a constant value of *ω* = 0.071 s^−1^ [12]. To the best of our knowledge, the dependence of this perfusion rate on the radius of CT-visible vessels has not been studied.

To motivate the present work, simulations comparing the perfusion-based model for large blood vessels to the advection-diffusion model have been conducted. The simulation geometry consisted of the mono-polar Angiodinamics Uniblate RF needle inserted into healthy liver tissue with a single cylindrical blood vessel placed parallel to the RF needle at a wall-to-wall distance of 1 mm. The RF needle active length was 1.5 cm. For the perfusion-based model the value *ω* = 0.05 s^−1^ was used inside the visible blood vessel. Distributed blood perfusion, mimicking fine-scale vessels was also used in both simulations (perfusion-based and advection-based) with the following temperature dependent blood perfusion rate

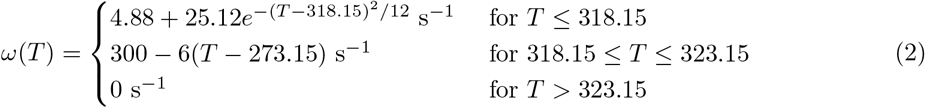

Figure 1 shows the thermal lesions obtained from these simulations. The radius values involved were 0.5 mm, 3 mm, and 5 mm. For the radius of 0.5 mm the perfusion based model substantially overestimated the thermal lesion. The perfusion-based lesions were much closer to the advection-based one for the other two radii but still overestimated the lesion extent. The relative errors in the perfusion-based lesion volumes (|*V*_perf_ − *V*_adv_|/*V*_adv_) are 72%, 73.6%, and 74.5% for *r* = 0.5 mm, *r* = 3 mm, and *r* = 5 mm respectively. This demonstrates the need to investigate the dependence of *ω* on *r*.

**Fig. 1.**
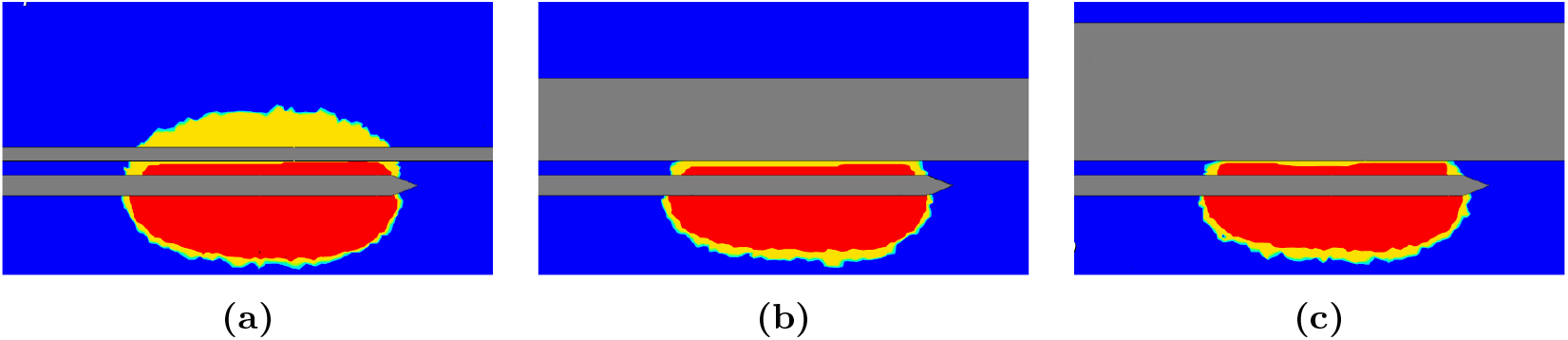
Comparison between the modified Pennes perfusion model of Kröger et al. and the advection diffusion model for blood vessel radii of: **(a)** 0.5 mm, **(b)** 3 mm, **(c)** 5 mm. The RF needle active length was 1.5 cm. The red region shows the extent of the thermal lesion from the advection-diffusion based model, while the yellow region shows the extent of the thermal lesion from the perfusion-based model. The blue colour indicates viable tissue.

The objective of the present work is to investigate, for a hepatic vessel radius *r* (ranging from 0.55 mm to 5 mm), the existence of a ‘best-estimate’ perfusion rate *ω_b_(r)* (subscript *b* indicates that it is the best estimate for that radius) such that the cooling effect of the perfusion-based model matches that of the advection-based model as closely as possible.

## 2 Methods

In the present work, simulations of RFA containing a single cylindrical blood vessel were performed to estimate the value of the perfusion rate as a function of vessel radius. The advection-diffusion model was assumed to be the exact model of the blood vessel and was used to determine the ‘truth’ thermal lesion. In the approximate model, the advection term is to be replaced with the perfusion term, but the best value of the perfusion rate is not known. The best perfusion rate for any radius can be defined as the one that minimizes the error in the predicted thermal lesion. The existence of such a perfusion rate has been investigated by setting up a PDE-constrained optimization problem.

### 2.1 Simulation Geometry

The computational geometry (Ω), shown in Figure 2a, consists of a cubic tissue region (Ω_*t*_) of side 10 cm, a blood vessel (Ω_*b*_) of variable radius *r*, and a simplified Angiodynamics UniBlate RF needle (diameter 1.5 mm) (Ω_*n*_) at a wall-to-wall distance of 1 mm from the vessel. The active length of the RF-needle was set to 20 mm. The internal structure of the RF needle is not homogeneous. To reduce the computational complexity the RF needle was assumed to be homogeneous and volume-averaged thermal and electrical properties were used in its interior. During the RFA procedure the current passed through the active surface of the needle (Figure 2b), and towards the ground pad, which corresponds to the bottom surface of the tissue cube.

**Fig. 2.**
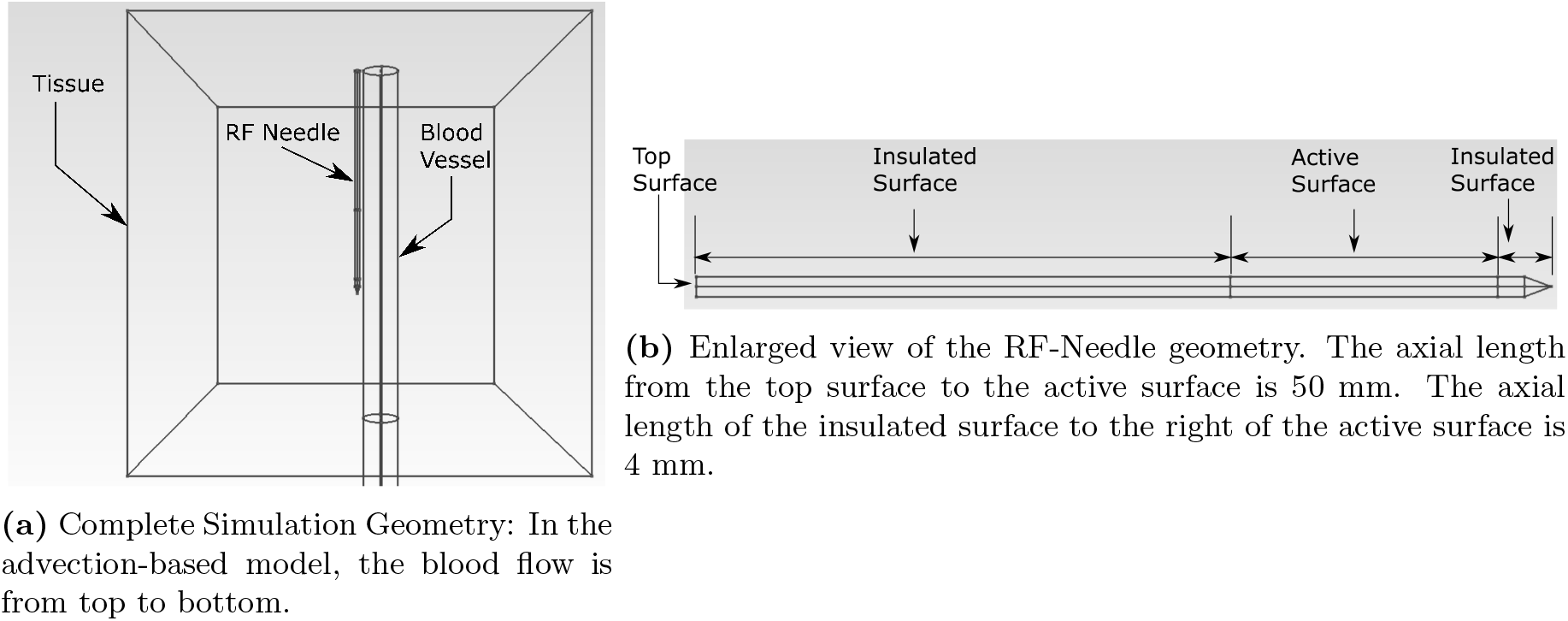
Geometry used for the simulations

### 2.2 Governing Equations

As in our previous work [9], the focus of the present work is on large vessel cooling effects. Hence the effect of fine-scale blood vessels was ignored in both advection-diffusion and perfusion-diffusion models.

#### 2.2.1 Advection-based Model

In the advection-based model the governing equations for solid tissue, blood, and RF-Needle were given by

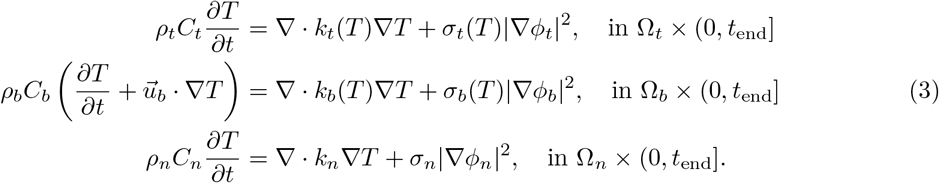

where the subscripts *t*, *b*, and *n* stand for tissue, blood, and RF needle respectively. The transient simulation was run for a total physical time of *t*_end_ = 540 s, which is the typical RFA treatment time used in clinical practice. The temperature and potential in the various subdomains is *T* and *ϕ*. Here *ρ_i_*, *C_i_*, *k_i_*, *σ_i_*, and *σ_i_*|∇*ϕ*|^2^ are the density, specific heat, thermal conductivity, electrical conductivity, electric potential, and the RF heat source density for *i* ∈ {*t, b, n*}. The term 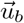 is responsible for heat advection, where 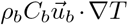 is the blood velocity. The blood flow in the vessel was assumed laminar and steady, leading to a Poiseuille flow profile for a perfectly cylindrical vessel [13]. Murray’s Law, a heuristic relationship between fluid flow-rate and channel diameter in biological systems, was used to determine the blood flow rate as a function of the vessel radius [14]. Murray’s Law is given by

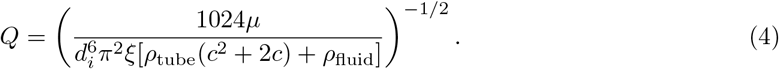

Here *d_i_*, *Q*, *μ*, *ξ*, and *c* are the vessel diameter, volume flow-rate, dynamic viscosity of the fluid, power-to-mass ratio of the pumping system, and a constant material property of the tube, respectively. The constants *ρ*_tube_ and *ρ*_fluid_ are the tube material density and fluid density respectively. The material constant ξ[*ρ*_tube_(*c*^2^ + 2*c*) + *ρ*_fluid_] is independent of the vessel diameter, and its value was derived from Eq. 4 by plugging in the known values of *Q* = 1.57e-5 m^3^·s^−1^ and μ = 0.00365 Pa·s for hepatic vessels with d_i_ = 10 mm [15]. The constant ξ[ρ_tube_(c^2^ + 2c) + *ρ*_fluid_] was estimated at 93.44 kg·s^−3^ · m^−1^. The electric potential was assumed to be governed by the quasi-static approximation

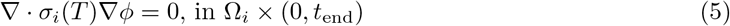

where *i* ∈ {*t, b, n*} [4].

The material property values used are listed in Table 1. Temperature dependent electrical and thermal conductivities were used for the blood and tissue subdomains. All specific heats and densities were assumed to be constant.

**Table 1.** Material property values used

| Region | Material property | Value |
| --- | --- | --- |
| Tissue | $\rho \text{ kg}\cdot\text{m}^{-3}$ | 1079 [16] |
| | $C \text{ J}\cdot\text{kg}^{-1}\cdot\text{K}^{-1}$ | 3540 [16] |
| | $k(T \text{ }^\circ\text{C}) \text{ W}\cdot\text{m}^{-1}\cdot\text{K}^{-1}$ | $4.69\text{e} - 1 + 1.16\text{e} - 3 \times T$ [17] |
| | $\sigma(T \text{ }^\circ\text{C}) \text{ S}\cdot\text{m}^{-1}$ | $-1.26\text{e} - 7 \times T^4 + 2.57\text{e} - 5 \times T^3 - 1.84\text{e} - 3 \times T^2$<br>$+ 6.35\text{e} - 2 \times T - 4.79\text{e} - 1$ [18] |
| | $A \text{ s}^{-1}$ | 3.1e98 [19] |
| | $\Delta E_a \text{ J}$ | 6.285e5 [19] |
| | $R_{\text{IG}} \text{ J}\cdot\text{mol}^{-1}\cdot\text{K}^{-1}$ | 8.314 [19] |
| Blood | $\rho \text{ kg}\cdot\text{m}^{-3}$ | 1050 [16] |
| | $C \text{ J}\cdot\text{kg}^{-1}\cdot\text{K}^{-1}$ | 3617 [16] |
| | $k(T \text{ }^\circ\text{C}) \text{ W}\cdot\text{m}^{-1}\cdot\text{K}^{-1}$ | $0.552 - 2.26\text{e} - 3 \times (37 - T)$ [20] |
| | $\sigma(T \text{ }^\circ\text{C}) \text{ S}\cdot\text{m}^{-1}$ | $100 \times (326.0 - 5.63 \times T)^{-1}$ [21] |
| RF<br>Needle | $\rho \text{ kg}\cdot\text{m}^{-3}$ | 1 |
| | $C \text{ J}\cdot\text{kg}^{-1} \text{ K}^{-1}$ | 1000 |
| | $k \text{ W}\cdot\text{m}^{-1}\cdot\text{K}^{-1}$ | 0.02 |
| | $\sigma \text{ S}\cdot\text{m}^{-1}$ | 1e7 |

The temperature and temperature gradient were continuous across all interfaces between the sub-domains. A convection boundary condition,

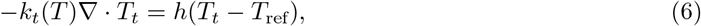

was applied to the tissue boundary, where *h* = 6 W ·m^−2^·K^−^1 is the convection coefficient and *T*_ref_ = 310 K is the ambient temperature. The top and side surfaces of the tissue boundary were electrically insulated, while the bottom surface (ground) had *ϕ* = 0 V. A time-dependent voltage

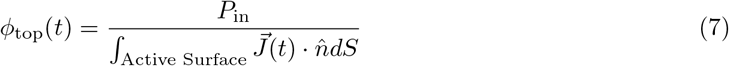

was applied at the top boundary of the RF-Needle, where 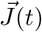 is the current density, and *P*_in_ = 6.024 W is the input power. All external boundaries of the blood-vessel domain were electrically insulated. The external curved boundary of the vessel was assumed to be thermally insulated, while the blood inlet and outlet boundaries were kept at 310 K. All tissue-vessel interfaces were assumed to be thermally and electrically conducting. All tissue-needle interfaces were assumed to be thermally conducting.

#### 2.2.2 Perfusion-based Model

In the perfusion-based model, the advection term 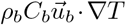 from Equation (3) was replaced with the perfusion term *ωρbCb(T − T_0_)*, where *ω* is the perfusion rate and *T*_0_ = 310 K. The energy conservation equation in the blood vessel subdomain becomes

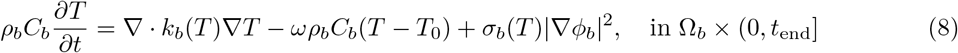

All other settings were identical to the advection-based model.

#### 2.2.3 Tissue Damage Model

The Arrhenius model was used to describe the time-evolution of tissue state [19]. Instead of the familiar Arrhenius damage index Ω, the tissue survival fraction Ψ = e^−Ω^ was used. The survival fraction Ψ obeys the following equation

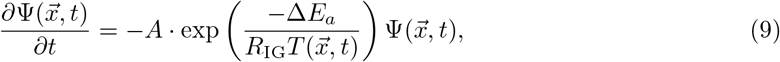

where *A* (s^−1^) is the frequency factor, ΔE*_a_* (J) is the activation energy, *R*_IG_ (J · mol^−1^·K^−1^) is is the ideal gas constant (values given in Table 1). The function 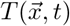 is the tissue temperature. The initial condition is 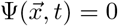, and the threshold for tissue necrosis is 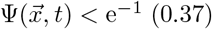. The final thermal damage field 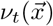 is defined as:

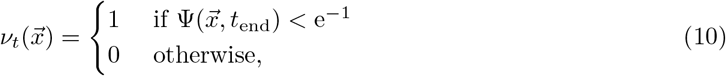

which assigns a value of 1 to the region corresponding to dead tissue, while 0 is assigned otherwise.

### 2.3 Non-dimensionalization

The governing equations were non-dimensionalized to improve the condition number of the mass- and stiffness-matrices.

#### 2.3.1 Advection-based Model

Non-dimensional variables and operators are notated with a ∗ super-script. The temperature *T*, electric potential *ϕ*, time *t*, position 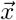, the del operator ∇, and the time derivative operator 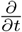 are non-dimensionalized in the following way:

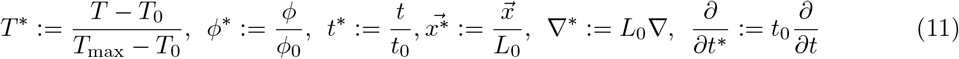

where *T*_0_ = 310.15 K, *T*_max_ = 373.15 K, *ϕ*_0_ = 25 V, *t*_0_ = *t*_end_ = 540 s, and *L*_0_ = 0.1 m. The following non-dimensional groups exist (for *i* ∈ {*t, b, n*}):

- Heat capacities 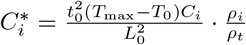
- Thermal conductivities 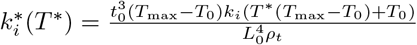
- Electrical conductivities 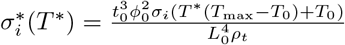
- Velocity 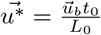

Using the above non-dimensional groups, the following non-dimensional form of the governing equations is obtained:

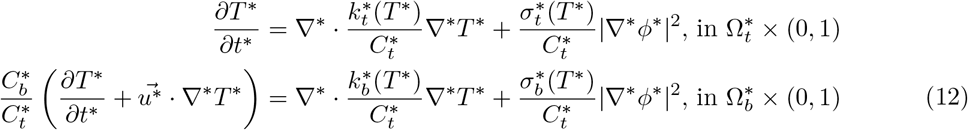

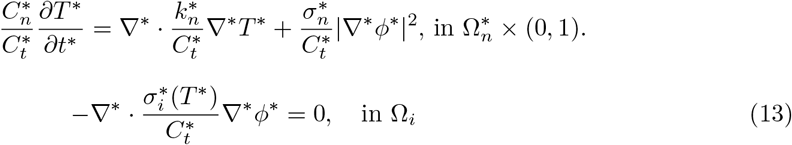

The convection boundary condition (from Equation (6)) in dimensionless form is:

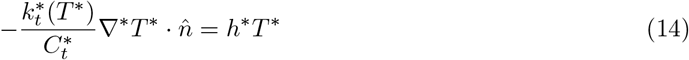

where the dimensionless convection coefficient is given by 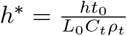. The electric potential boundary condition from Equation (7) in dimensionless form is

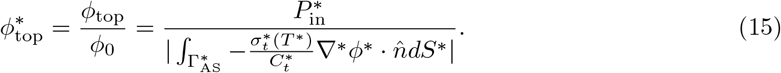

The dimensionless input power is given by 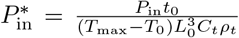. The dimensionless current *I*^∗^ is related to the dimensional current *I* by

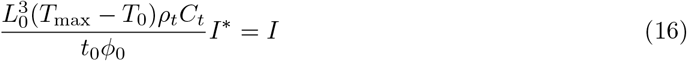

#### 2.3.2 Perfusion-based Model

The non-dimensionalization of the perfusion rate *ω* is

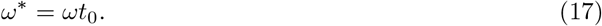

The other terms appearing in Equation (8) are non-dimensionalized the same as shown earlier. The non-dimensionalized form of the energy conservation equation for the blood vessel is given by:

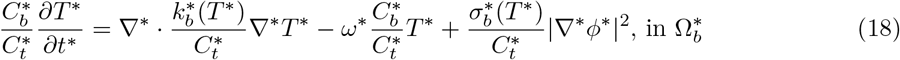

The dimensionless forms of the other equations are identical to the advection-based case.

### 2.4 Optimization Problem

The goal is to determine the function 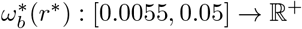 such that, for a given value of non-dimensional blood vessel radius *r*∗, the perfusion-based model (eq. (18)) using the value 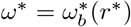 gives the best estimate of the thermal lesion when the advection-based result for the same vessel radius is assumed to be the truth. In an attempt to determine this best estimate, the following PDE-constrained optimization problem was set up:

For *r*^∗^ ∈[0.0055, 0.05] find 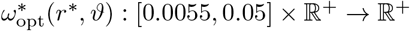 such that

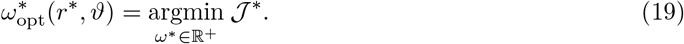

The objective functional 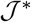 is defined as

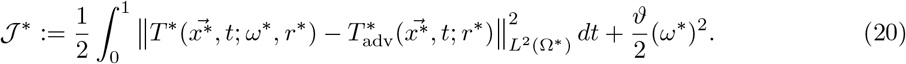

The temperature fields 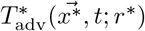 and 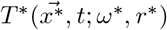 are solutions to problems (12) and (18) respectively with the radius of 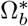 being 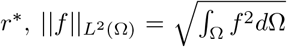 is the *L*^2^ norm, and *ϑ* ∈ ℝ^+^ is a regularization parameter. This choice of objective function was made to satisfy two constraints: (*i*) 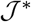 should be differentiable with respect to *ω*^∗^, and (*ii*) 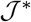 should become smaller as the thermal lesion predicted by the perfusion-based model approaches the one predicted by the advection-based model. The accuracy of an optimal solution to Problem (19), 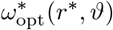, was evaluated using the following relative error

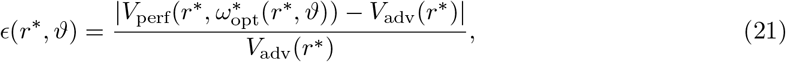

where 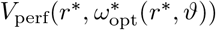 is the thermal lesion obtained from the perfusion-based model with vessel radius *r*^∗^, using the perfusion rate *ω*_opt_(*r, ϑ*), and *V*_adv_(*r*^∗^) is the thermal lesion obtained from the advection based model, for the same vessel radius. For any *r*, the best estimate 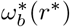 was defined as:

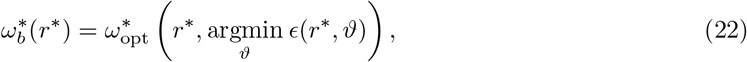

where the minimization of the relative error *ϵ*(*r*^∗^, *ϑ*) is performed over all the *ϑ* values considered.

### 2.5 Numerical Implementation

The finite element method-based software library MOOSE framework was used for the implementation [22]. The SUPG stabilization scheme was used to compute the weak form of the advection term [23]. This is necessary when FEM is used for advection-dominated problems. The Additive Schwarz Method (ASM) pre-conditioner was used for the linear solves which are part of Newton’s method for non-linear solves. The ASM pre-conditioner is ideal for non-linear PDEs which are to be solved in parallel. It involves decomposition of the domain into overlapping subdomains. The original problem is solved on the individual subdomains and the solutions are then combined. For each of the subdomain solves, the incomplete LU-decomposition pre-conditioner was chosen. MOOSE Framework makes use of the PETSc library for the linear algebra computations, and it contains implementations of both ASM and LU-decomposition pre-conditioners [24].

A tetrahedral mesh was generated using the 3D finite element mesh generation tool Gmsh [25]. A mesh convergence study was performed to determine the appropriate mesh-size (which led to around 17,000 tetrahedral elements) and a time-step of 0.5. The backward Euler method was used to discretize the time-derivatives.

The optimization routine was run for the following 10 (non-dimensionalized) radius values lying in 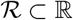

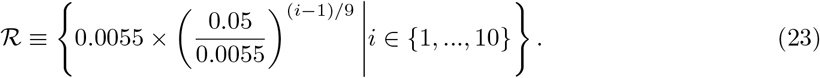

The Fletcher-Reeves Conjugate Gradient (FRCG) Method was used to optimize the objective function [26]. The optimization library DAKOTA was used to solve the optimization problem [27]. The termination condition for the optimization algorithm was a threshold of 1e-4 for the relative change in the value of the objective function between successive iterations. The FRCG method requires the gradient of the objective function with respect to the design variables. This derivative was computed by solving the tangent linear system of the original forward problem (Eq. (18)) [28]. The tangent linear system was solved at each time-step after the Newton solve for that time-step. This made it possible to compute the objective function value and its gradient in a single forward solve albeit with a slightly increased time-requirement. This was also done using the MOOSE framework.

The optimal value of the regularization parameter *ϑ* in the objective function (Equation (20)) is not known *a priori*. The purpose of the regularization parameter is to ensure that the objective function 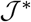 is convex. If the objective function 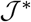 is convex for a particular value of the regularization parameter, the optimum 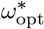 is unique and hence independent of the initial guess. An increasing value of *ϑ* puts a stronger penalty on a high value of the perfusion rate 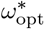, thus making the objective convex, but it also causes the regularization term to dominate over the temperature difference term. Thus, *ϑ* needs to be tuned to ensure the convexity of 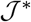, while at the same time maintaining the relative importance of the temperature difference term. The value of theta is expected to influence the optimal value since it changes the properties of the objective function. To determine the range of suitable values, multiple optimizations were performed with a very coarse mesh with different values of the regularization parameter *ϑ* and different initial guesses for *ω*^∗^. For *ϑ* ∈ {5e-8, 1e-8, 5e-9, 2e-9, 1e-9, 1e-10}, the initial values 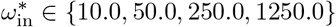 were tested for all 10 radius values. Based on the results of these coarse-mesh simulations, a few most appropriate values of *ϑ* were selected for the fine-mesh calculations.

## 3 Results

### 3.1 Effect of Varying Regularization Parameter

For the coarse-mesh calculations, for each radius and *ϑ* value the mean and standard deviation of 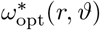 was computed for all 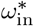 values. These results are shown in Figure 3a. The standard deviation of 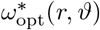 is quite high for *ϑ* = 1e-10 for r > 2 mm. For *ϑ* = 1e 9 there is some instability in 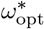 for r = 3.91 mm and r = 5 mm. For the rest of the *ϑ* and r values, the objective function admits a more stable optimum. The smallest values of *ϑ* that admit a stable optimum are the best to maintain the relative importance of the temperature difference term.

**Fig. 3.**
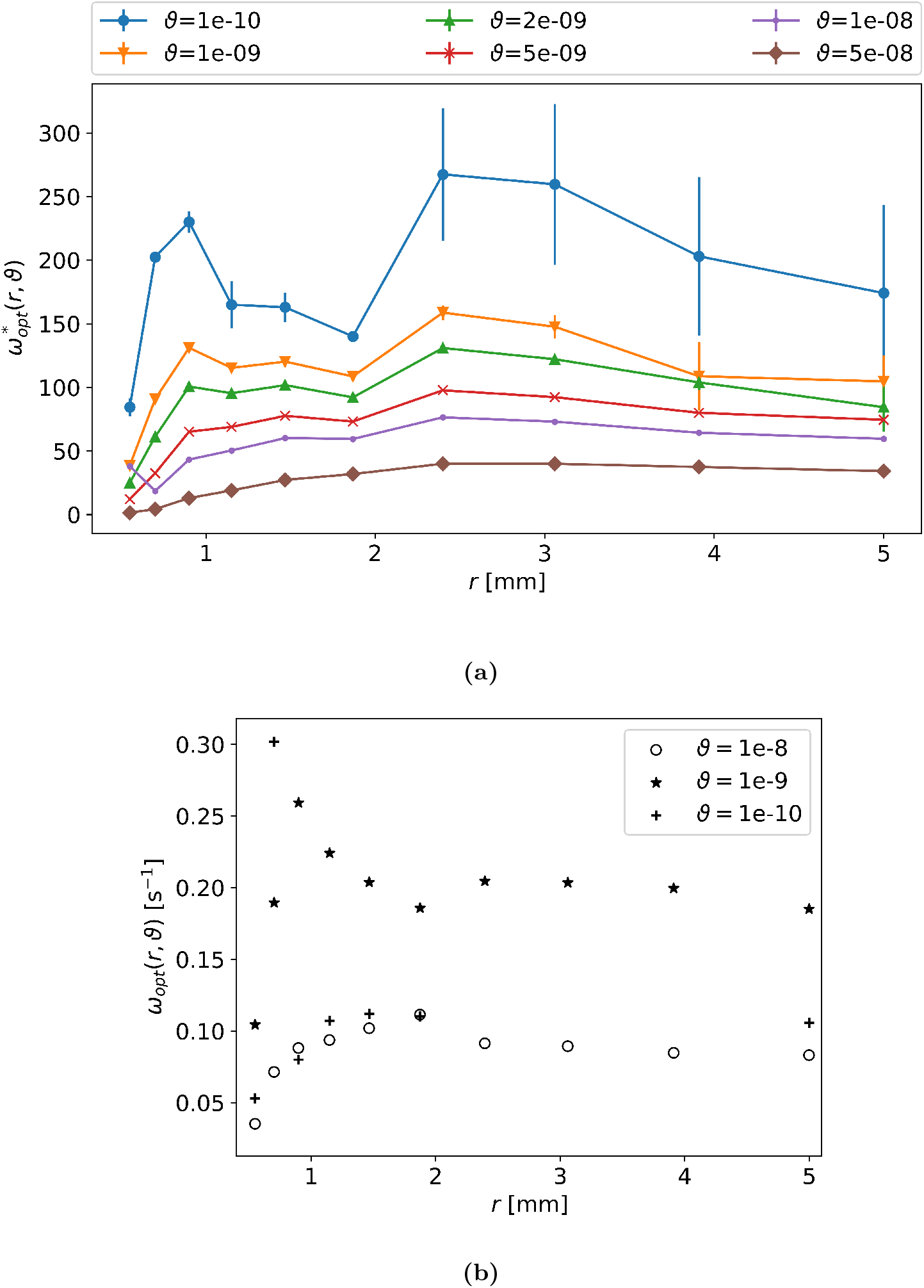
(a)Results of variation in 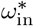 on the optimum 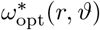 for different radii *r* and and different values of the regularization parameter *ϑ* using a coarse mesh. **(b)** variation in the optimal perfusion rate *ω*_opt_(*r*, *ϑ*) [s^−1^] (in dimensional form) with vessel radius r [mm] for all values of the regularization parameter *ϑ* considered with a fine mesh.

Based on these results the values *ϑ* = 1e − 10, *ϑ* = 1e − 9, and *ϑ* = 1e − 8 were chosen for the fine-mesh calculations. Figure 3b shows the optimal values *ω*_opt_(*r*, *ϑ*) found for the different values of *r* and *ϑ* considered. The optimization was first performed for all *r* using *ϑ* = 1e-8 and *ϑ* = 1e-9. For *ϑ* = 1e-8 *ω*_opt_(*r*) first increases with increasing radius, reaches a maximum for r = 1.87 mm and then decreases for r = 2.40 mm. The value of *ω*_opt_(r, *ϑ*) does not change much with increasing radius from that point onwards. For *ϑ* = 1e-9, *ω*_opt_(*r*, *ϑ*) first increases with increasing *r* and reaches a maximum at *r* = 0.90 mm. Then it oscillates slightly with increasing *r*. For *r* = 2.4 mm, *r* = 3.06 mm, and *r* = 3.91 mm the thermal lesion volumes obtained using the perfusion-based model with *ω*_opt_(*r*, *ϑ* = 1e-9) are within 3% of the corresponding advection-based lesion volumes (see Table 2). This is sufficient accuracy, and for these radii the best estimate for the perfusion rate, *ω*_b_, has been found. For the remaining radii neither *ϑ* = 1e-8 nor *ϑ* = 1e-9 results in optimum *ω*_opt_(*r*, *ϑ*) values which lead to acceptable relative errors in the perfusion-based lesion volume. In an attempt to address this issue, the optimization procedure was repeated with *ϑ* = 1e-10 for *r* ∈ {0.5 mm, 0.7 mm, 0.9 mm, 1.15 mm, 1.47 mm, 1.87 mm, 5 mm}. These results are also in Fig. 3b. For *ϑ* =1e-10 the value of *ω*_opt_(*r*, *ϑ*) increases with increasing radius and reaches a plateau of 0.1 s^−1^ with increasing radius, with a large spike for *r* = 0.7 mm.

**Table 2.** Relative error between *V*_perf_ and *V*_adv_ for all *r* and *ϑ* considered: The *ϵ*_min_ column shows the minimum value of relative error over all values of *ϑ* considered for each radius. The final column shows the best estimate, *ω*_b_, for the perfusion rate value obtained from the optimization using different values of *ϑ*.

| $r \text{ [mm]}$ | $V_{\text{adv}} \text{ [cm}^3\text{]}$ | $\epsilon[\%]$ | | | $\epsilon_{\text{min}}[\%]$ | $\omega_b(r) \text{ [s}^{-1}\text{]}$ |
| --- | --- | --- | --- | --- | --- | --- |
| | | $\vartheta = 1\text{e-}8$ | $\vartheta = 1\text{e-}9$ | $\vartheta = 1\text{e-}10$ | | |
| 0.55 | 4.21 | 26.34 | 21.73 | 24.93 | 21.73 | 0.105 |
| 0.70 | 3.72 | 26.34 | 24.24 | 19.43 | 19.43 | 0.302 |
| 0.90 | 3.46 | 34.68 | 23.04 | 35.71 | 23.04 | 0.259 |
| 1.15 | 3.17 | 36.94 | 20.27 | 32.78 | 20.27 | 0.224 |
| 1.47 | 3.11 | 21.10 | 9.12 | 18.81 | 9.12 | 0.204 |
| 1.87 | 3.37 | 10.15 | 5.45 | 10.27 | 5.45 | 0.186 |
| 2.40 | 3.28 | 8.40 | 1.71 | - | 1.71 | 0.205 |
| 3.06 | 3.02 | 9.54 | 2.88 | - | 2.88 | 0.203 |
| 3.91 | 2.90 | 7.55 | 0.72 | - | 0.72 | 0.196 |
| 5.00 | 2.71 | 5.48 | 4.37 | 2.48 | 2.48 | 0.105 |

### 3.2 Comparison of advection-based and perfusion-based Models

The *ω*_opt_(*r, ϑ*) values (which solve the optimization problem from Eq. (19)) shown in the previous section were used in forward simulations of RFA to determine perfusion-based estimates to the thermal lesion. These were compared to the advection-based estimates, which were treated as the ground truth. Figure 4 shows a comparison of the perfusion-based and advection-based lesions for *ϑ* = 1e-8, *ϑ* = 1e-9, and *ϑ* = 1e-10 respectively. For *r* = 0.55 mm (Fig. 4a) all perfusion-based thermal lesion estimates are larger than the truth. The perfusion based lesion is smallest for *ϑ*=1e-9. Also, for *r* = 0.70 mm (Fig. 4b) mm all perfusion-based thermal lesion estimates are larger than the ground truth. The perfusion-based lesion grows in size as *ϑ* increases. For both aforementioned values of radius, all lesion estimates envelop the vessel. For *r* = 0.90 mm and *r* = 1.15 mm (Fig. 4c and 4d) the perfusion-based lesion estimates are again larger than the advection-based ones. However, the perfusion-based volumes are smallest for *ϑ* =1e-9. All perfusion-based lesions completely surround the vessels, whereas the advection-based lesion does not. For *r* = 1.47 mm (Fig. 4e) The perfusion-based lesions are only slightly larger than the advection-based one. The perfusion-based lesion for *ϑ* =1e-9 is the smallest among all *ϑ* values and does not envelop the vessel (just like the advection-based lesion). For the other values of *ϑ* the perfusion-based lesion does envelop the vessel. For the rest of the radius values (Fig. 4f to 4j) the perfusion-based lesions appear almost identical to the advection-based lesions.

**Fig. 4.**
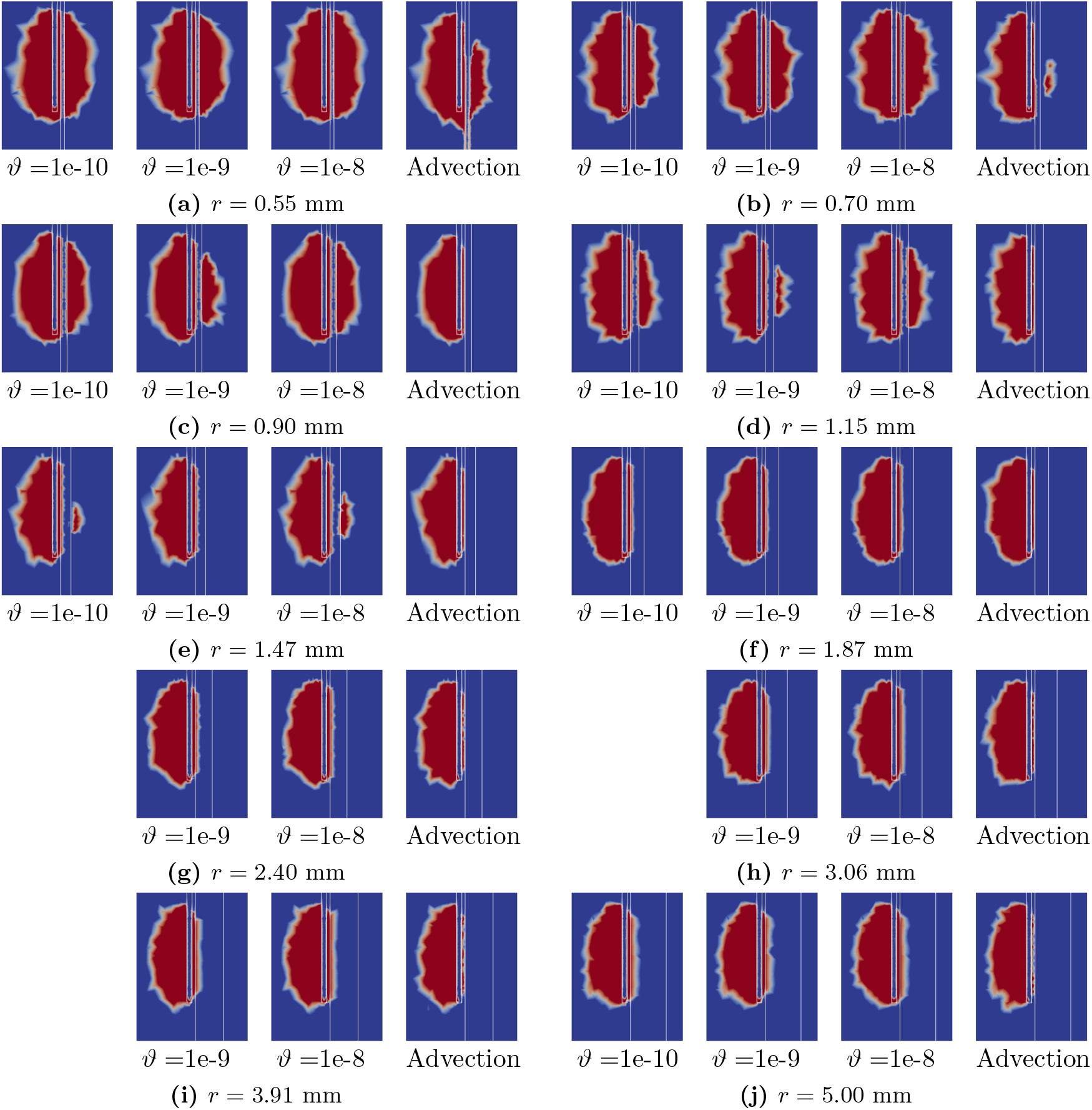
Comparison of thermal lesions obtained from the perfusion-based model (using *ω*_opt_(*r*, *ϑ*) for all values of *r* and *ϑ* considered) and then advection-based model: Sections through the vessel and RF needle axes are presented. The outlines of the RF needle and blood vessel are represented by white lines. The RF needle is located to the left of the blood vessel.

### 3.3 Best Estimates of the Perfusion Rate

A plot of the relative error *ϵ*(*r, ϑ*) (defined in E q. (21)) vs *r* for different values of *ϑ* (for the fine-mesh calculations) is shown in Fig. 5a. For all values of r the relative error in lesion volumes is smaller for *ϑ* = 1e-9 than for *ϑ* = 1e-8. For *ϑ* = 1e-10 the relative error is smaller than that for the remaining *ϑ* values for *r* = 0.7 mm and *r* = 5 mm. For the remaining values of radius the relative error for *ϑ* = 1e-9 is smallest over all *ϑ*. Table 2 shows the quantitative data for the relative error in thermal lesion volumes. The best estimates to the perfusion rate, *ω*_b_, for all 10 radii (determined according to Eq. 22) are also shown in that table. It is desirable to have *ω*_b_ [s^−1^] as a continuous function of the vessel radius *r* [mm]. Hence, Akima cubic splines were used to obtain such a function [29]. It is given by

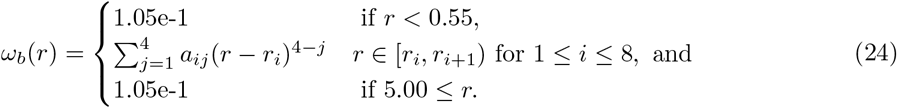

**Fig. 5.**
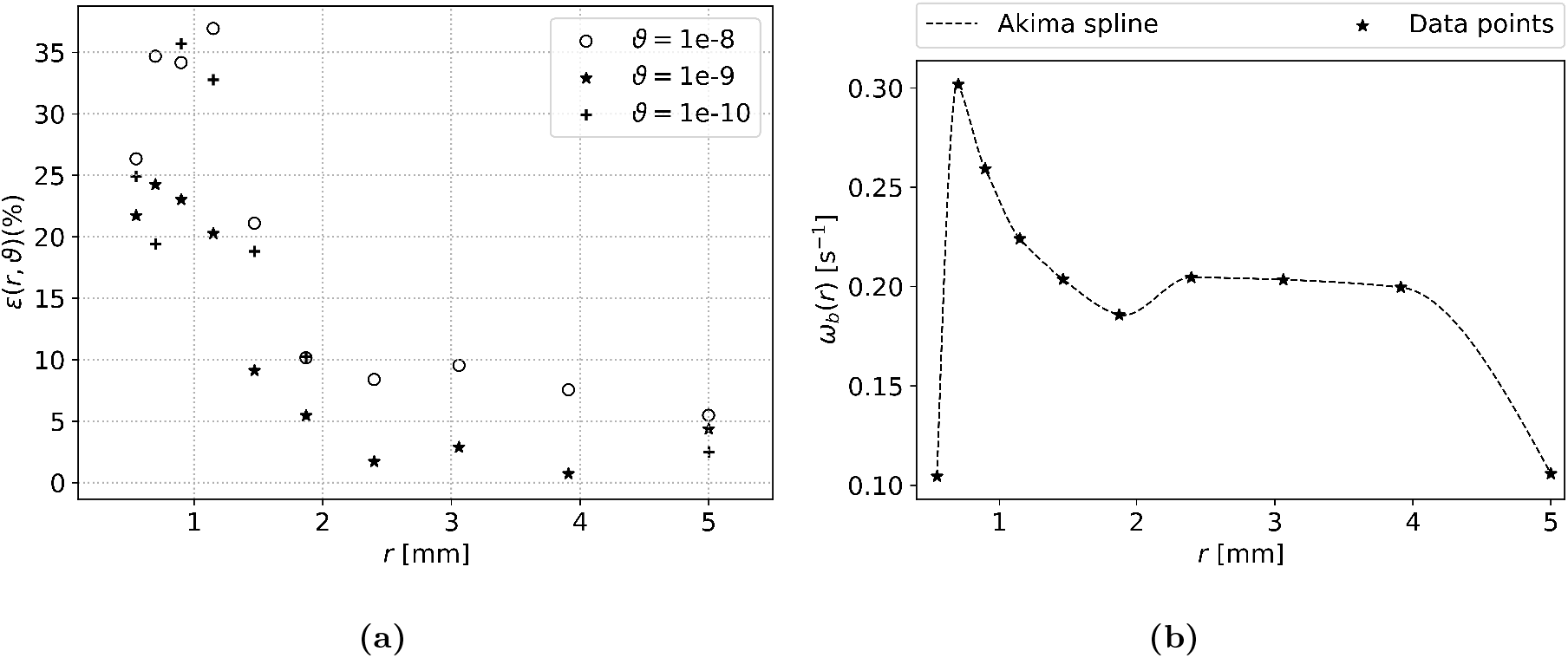
The plot shows **(a)** variation of the relative error ϵ(*r*, *ϑ*) (defined in Eq. (21)) in the thermal lesion obtained using the perfusion based model for all radius and regularization parameter values; **(b)** variation in the best estimate to the perfusion rate, *ω*_b_(*r*) [s^−1^], with vessel radius r in dimensional form. For any *r* the best value of the perfusion rate is the one for which the relative error in lesion volumes ϵ(*r*, *ϑ*) is smallest over all all *ϑ* considered.

The real coefficients *a_i_, b_i_, c_i_*, and *d_i_* for *i* = 1, …, 9 and the radii *r_i_* are given in Table 3. This continuous function along with the data points is shown in Fig. 5b.

**Table 3.**
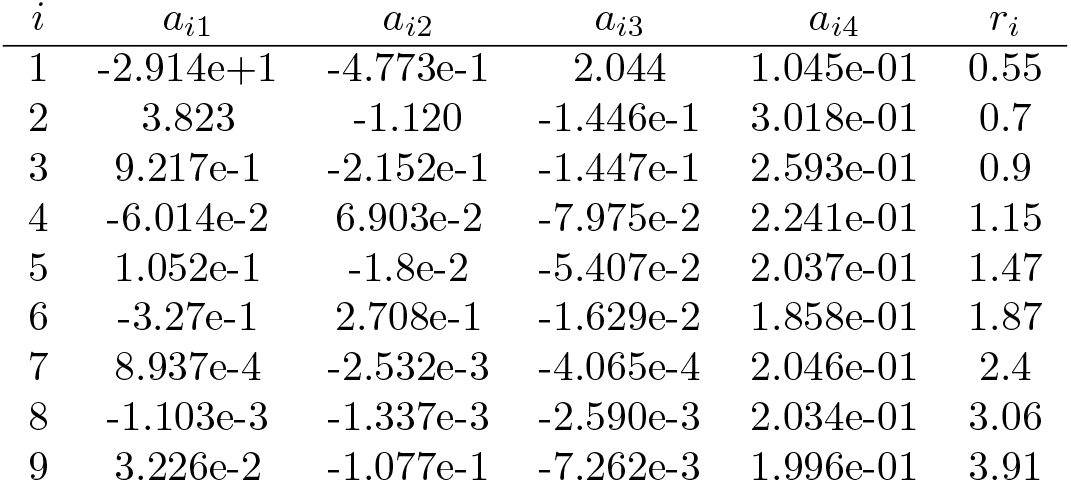
The table lists the Akima cubic spline coefficients *a_ij_* for *i* = 1, …, 9 and *r_i_* and *j* = 1, …, 4 from Equation (24) which gives the dependence of the best estimate of the perfusion rate *ω*_b_ [s^−1^] on *r* [mm].

| $i$ | $a_{i1}$ | $a_{i2}$ | $a_{i3}$ | $a_{i4}$ | $r_i$ |
| --- | --- | --- | --- | --- | --- |
| 1 | -2.914e+1 | -4.773e-1 | 2.044 | 1.045e-01 | 0.55 |
| 2 | 3.823 | -1.120 | -1.446e-1 | 3.018e-01 | 0.7 |
| 3 | 9.217e-1 | -2.152e-1 | -1.447e-1 | 2.593e-01 | 0.9 |
| 4 | -6.014e-2 | 6.903e-2 | -7.975e-2 | 2.241e-01 | 1.15 |
| 5 | 1.052e-1 | -1.8e-2 | -5.407e-2 | 2.037e-01 | 1.47 |
| 6 | -3.27e-1 | 2.708e-1 | -1.629e-2 | 1.858e-01 | 1.87 |
| 7 | 8.937e-4 | -2.532e-3 | -4.065e-4 | 2.046e-01 | 2.4 |
| 8 | -1.103e-3 | -1.337e-3 | -2.590e-3 | 2.034e-01 | 3.06 |
| 9 | 3.226e-2 | -1.077e-1 | -7.262e-3 | 1.996e-01 | 3.91 |

## 4 Discussion

Barring the *r* = 5 mm and *r* = 0.55 mm cases the best estimate of the perfusion rate *ω*_b_ generally decreases with increasing radius. In the perfusion based model, the magnitude of the blood vessel heat sink depends on the perfusion rate, and the volume of the vessel subdomain. If the same amount of heat is to be removed, a smaller vessel would need a larger perfusion rate, than a vessel of larger radius owing to the difference in their volumes. In the advection-based model, the heat sink effect depends on the flow-rate (given by Murray’s Law) and the vessel volume. Hence it decreases with decreasing radius, but its impact on the best estimate of the perfusion rate is not straightforward to predict. It is likely that the increase in the best estimate perfusion rate with decreasing vessel radius is due to the smaller volume of the vessel subdomain. This does not, however, explain the behaviour of the best estimate perfusion rate for *r* = 0.55 mm and *r* = 5 mm.

Recall that the objective function 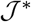 (see Eq. (20)) has a temperature difference term and a regularization term. As stated before, the temperature difference term is used as a proxy for the difference in the thermal lesions. For smaller radii the advection-based temperature field shows some heating in the neighbourhood of the blood vessel downstream of the RF needle (see Figures 6 and 7). It is possible that the optimization algorithm tries to match this temperature by keeping the *ω* value low, thereby explaining the small *ω*_b_ value for *r* = 0.55 mm. This directional effect of blood flow grows weaker with increasing radius [9]. The choice of the objective function could be improved, for example, by considering only the temperatures within a small region around the RF needle. This objective function may be able to capture the difference in the thermal lesions more accurately while still remaining differentiable with respect to *ω*. Figure 8 shows a plot of the dimensionless objective function 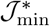, and its constituent terms: 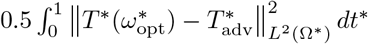 (temperature difference term) and 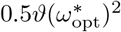 (penalty term) vs the radius r for all *ϑ* values considered. For all radii except 5 mm, the best estimate of the perfusion rate *ω*_b_ is the one with the smallest value of the temperature difference term (triangle markers in Fig. 8). The same trend is found for the value of the objective function (circle markers in Fig. 8). Decreasing *ϑ* from 1e-8 to 1e-9 (blue and green markers, respectively, in Fig. 8) results in a smaller value of the total objective function for all radii. This is due to a decrease in the value of both its constituent terms. The decrease in value of the temperature difference term indicates a closer match between the advection-based and perfusion-based temperature fields. When the value of *ϑ* is decreased further to 1e-10, the regularization term becomes extremely small, while the value of the temperature difference term increases in all cases except *r* = 0.7 mm. It is likely that the objective function is no longer convex and has some local minima for *ϑ* = 1e-10 (indicated by Fig. 3a), hence the optimization procedure terminates at a non-optimal value of *ω*^∗^. For *r* = 5 mm and *ϑ* = 1e-10 the temperature difference term is not the smallest, yet it gives the best estimate *ω*_b_ in terms of the thermal lesion relative error. This result is surprising, and it was investigated further. Figure 9 shows a plot of perfusion-based lesion volume versus *ω* for *r* = 5 mm and the associated relative error. Contrary to the expectation that the perfusion-based lesion volume would decrease monotonously as *ω* increases, there exists a local minimum. The reason for this behaviour is unclear, but could be the interaction between the constant power input and the temperature dependence of tissue electrical conductivity. The total current at the end of the simulation was found to decrease with increasing *ω*.

**Fig. 6.**
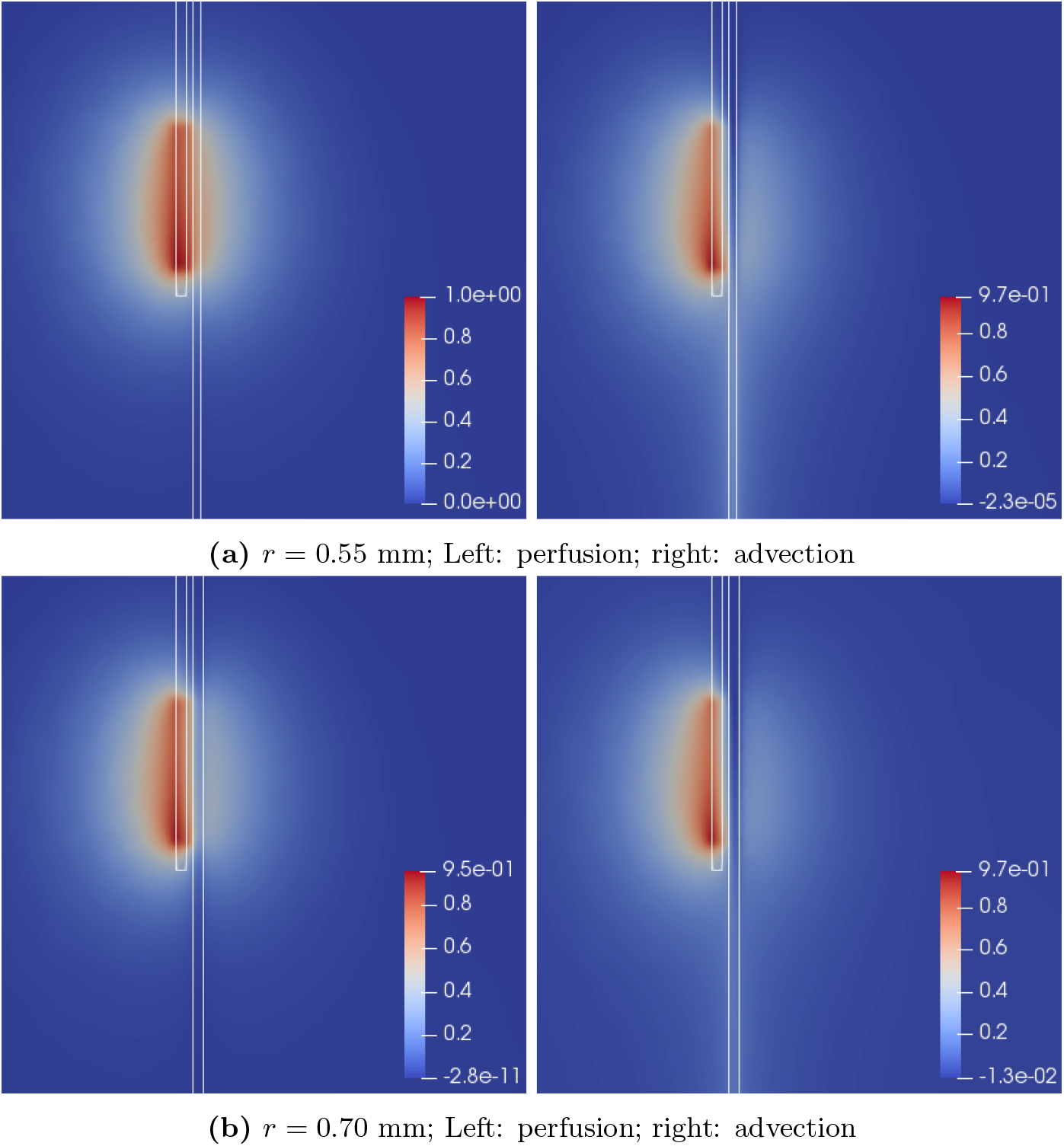
Comparison of final non-dimensional advection-based and perfusionn based temperature fields for *r* = 0.9 mm, *r* = 1.15 mm. The perfusion based temperature was computed using the best estimate perfusion rate *ω*_b_. In the advection-based temperature field heating around the vessel downstream of the RF needle decreases with increasing radius.

**Fig. 7.**
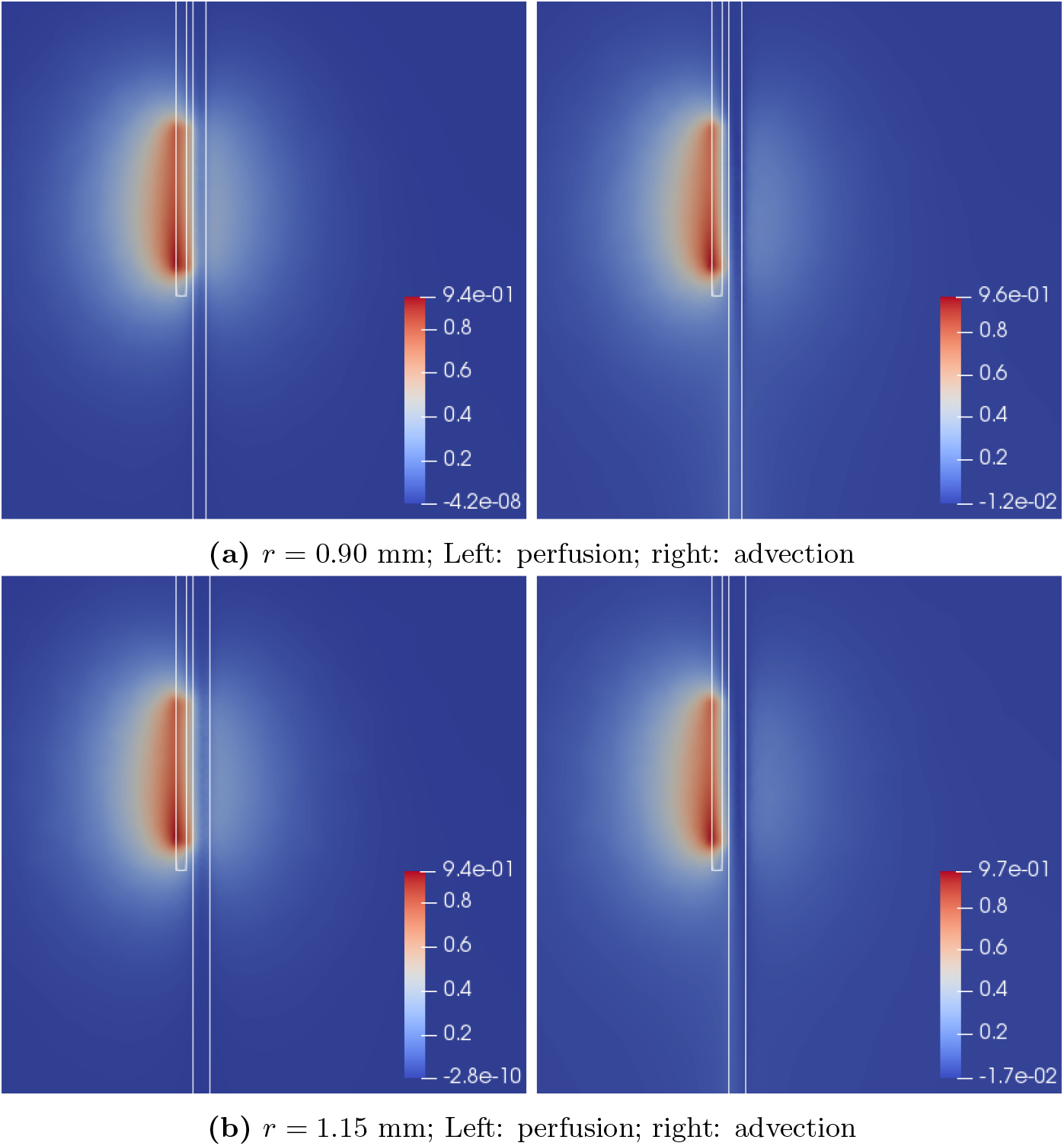
Comparison of final non-dimensional advection-based and perfusion based temperature fields for *r* = 0.9 mm, *r* = 1.15 mm. The perfusion based temperature was computed using the best estimate perfusion rate *ω*_b_. In the advection-based temperature field heating around the vessel downstream of the RF needle decreases with increasing radius.

**Fig. 8.**
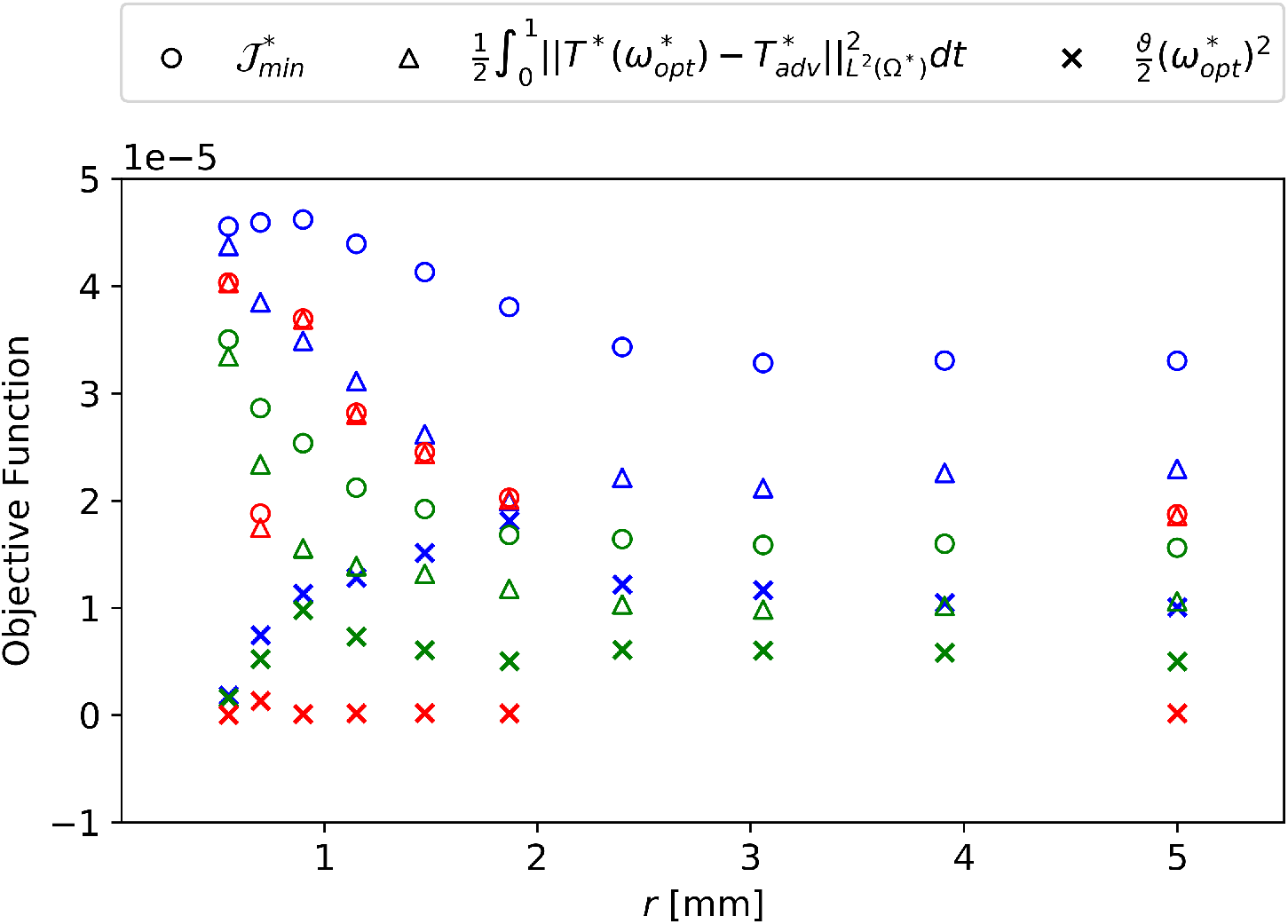
Plot of the final objective function 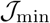, and its two constituent terms versus radius for all the *ϑ* values considered: The colour coding is as follows, Red: *ϑ* = 1e-10, Green: *ϑ* = 1e-9, Blue: *ϑ* = 1e-8.

**Fig. 9.**
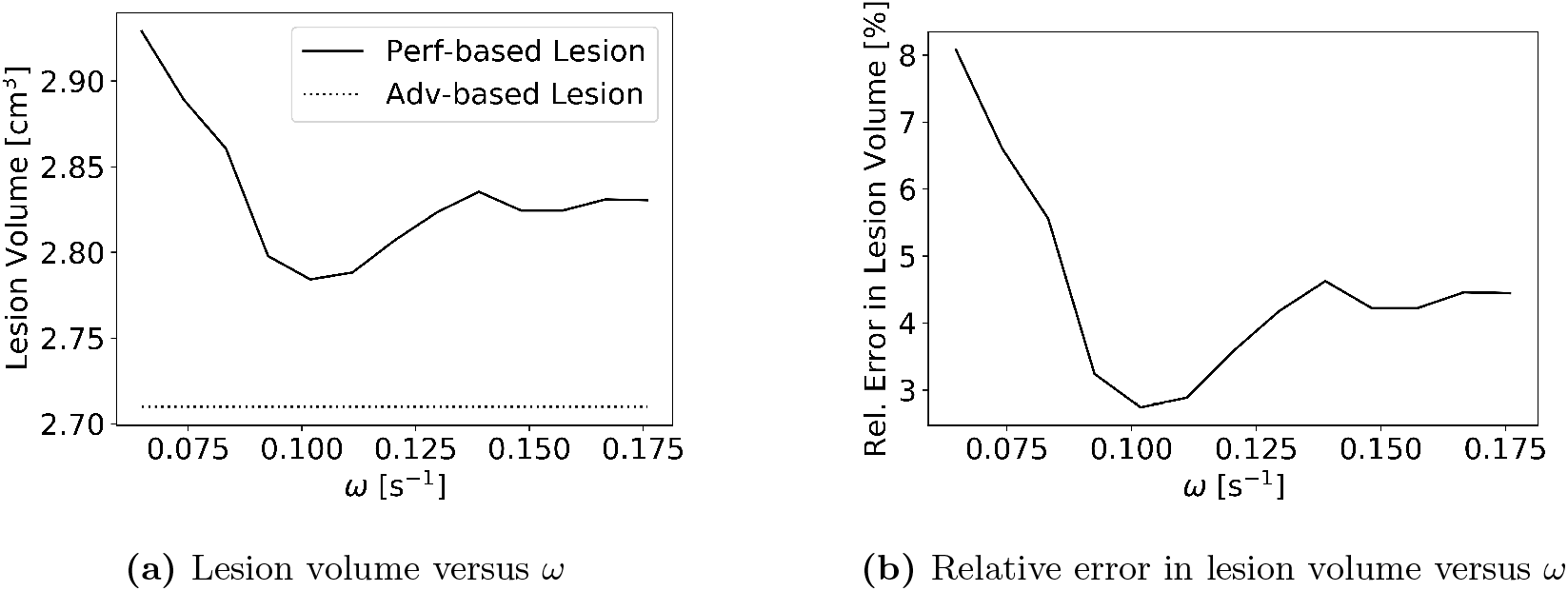
Effect of variation in the perfusion rate *ω* on the perfusion-based lesion volume and relative error for *r* = 5 mm: The relative error is computed assuming the advection based lesion as the truth.

Among the best perfusion rate estimates *ω*_b_ for the 10 values of radius shown in Fig. 5b, four lead to thermal lesions of acceptable accuracy: 2.4 mm, 3.06 mm, 3.19 mm, and 5 mm. For the remaining radii, i.e. 0.55 mm ≤ *r* ≤ 1.87 mm, the relative errors are rather high. This could have multiple reasons: (*i*) the form of the objective functions used is not the best for this application, (*ii*) for these radii, the perfusion-based model is incapable of matching the advection-based lesion more closely. Further investigation can focus on determining which of the above statements is true. One constraint in the choice of the objective function is its differentiability with respect to the perfusion rate *ω*. This property makes it possible to use a gradient-based optimization method, which saves computational effort. An objective function which involves the difference between thermal damage fields will not be differentiable with respect to *ω* and will warrant gradient-free optimization methods which typically require many more forward solves than gradient-based ones. A hybrid approach can be used which uses the gradient based estimate for *ω* as an initial guess for a gradient-free optimization using a tissue-damage-based objective function. The hybrid approach could help determine whether statement (*ii*) above is true. Additionally, repeating the procedure for more radius values in the range of interest could give more information about the behaviour of the optimal perfusion rate with changing radius.

The results presented in this chapter are dependent on the values of the geometrical parameters (i.e. vessel orientation, RF needle active length, wall-to-wall distance between needle and vessel). A generalization of the results obtained which considers the variation in the other parameters would make the approximate model more accurate. Another limitation of this work is that the blood flow rate is assumed to be governed by Murray’s law. Since Murray’s law is only a heuristic relationship, it would be useful to investigate the sensitivity of the results obtained to variations in blood flow rate for the range of vessel radii considered. Furthermore, idealized cylindrical blood vessels have been considered in the present work. In reality the vessels would have further complications such as curved axes, variable cross-sections and branching which introduces a lot of new parameters. Future work can focus on addressing these concerns.

## 5 Conclusion

In this work the possibility of using the Pennes model to predict large vessel cooling effects for 0.55 mm ≤ *r* ≤ 5 mm has been explored and optimal values for the perfusion rate have been identified. For 2.40 mm ≤ *r* ≤ 5 mm the perfusion-based model was found to approximate the advection-based model with a lesion volume relative error of under 3%. For *r* = 1.87 mm the relative error was 5.45%. For 0.55 mm ≤ *r* ≤ 1.47 mm the errors associated with the perfusion based model are ~10%. Suggestions have been made for future work to address the problem of low accuracy of the perfusion-based model in that range of radius values. Though large for the particular application, these errors are small compared to those obtained using perfusion rate values from literature. A continuous function *ω*_b_(*r*), which makes use of Akima splines, has been presented for the optimal perfusion rate which is valid for the range of radius values considered. This function can improve the accuracy of existing perfusion-based RFA models, which simplify the numerical implementation by removing the need for additional stabilization (e.g. SUPG). This simplification makes this class of models more suitable for use in applications where fast estimation of RFA outcomes is crucial (e.g. for real time intervention support).

### 6 Funding Information

This work was supported by the European Commission under the Marie Skłodowska Curie Actions - Innovative Training Networks (MSCA-ITN) grant of the type European Industrial Doctorate (EID) (project number 642445).

## List of Tables

1. Material property values used … … … … … … … … … … . .
2. Relative error between *V*_perf_ and *V*_adv_ for all *r* and *ϑ*. considered: The ε_min_ column shows the minimum value of relative error over all values of *ϑ* considered for each radius. The final column shows the best estimate, *ω_b_*, for the perfusion rate value obtained from the optimization using different values of *ϑ*. … … … … … … … …
3. The table lists the Akima cubic spline coefficients *a_ij_* for *i* = 1, …, 9 and *r_i_* and *j* = 1, …, 4 from Equation (24) which gives the dependence of the best estimate of the perfusion rate *ω_b_* [s^−1^] on *r* [mm]… … … … … … … … … … .

## List of Figures

1. Comparison between the modified Pennes perfusion model of Kroger et al. and the advection diffusion model for blood vessel radii of: (a) 0.5 mm, (b) 3 mm, (c) 5 mm. The RF needle active length was 1.5 cm. The red region shows the extent of the thermal lesion from the advection-diffusion based model, while the yellow region shows the extent of the thermal lesion from the perfusion-based model. The blue colour indicates viable tissue… … … … … … … … … … … … .
2. Geometry used for the simulations … … … … … … … … … . .
3. **(a)** Results of variation in 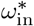 on the optimum 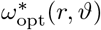 for different radii *r* and and different values of the regularization parameter *ϑ* using a coarse mesh. **(b)** variation in the optimal perfusion rate *ω*_opt_(*r*, *ϑ*) [s^−1^] (in dimensional form) with vessel radius *r* [mm] for all values of the regularization parameter *ϑ* considered with a fine mesh… .
4. Comparison of thermal lesions obtained from the perfusion-based model (using *ω*_opt_(*r*, *ϑ*) for all values of *r* and *ϑ* considered) and then advection-based model: Sections through the vessel and RF needle axes are presented. The outlines of the RF needle and blood vessel are represented by white lines. The RF needle is located to the left of the blood vessel… … … … … … … … … … … … … … … .
5. The plot shows **(a)** variation of the relative error ϵ(*r*, *ϑ*) (defined in Eq. (21)) in the thermal lesion obtained using the perfusion based model for all radius and regularization parameter values; **(b)** variation in the best estimate to the perfusion rate, *ω*_b_(*r*) [s^−1^], with vessel radius r in dimensional form. For any r the best value of the perfusion rate is the one for which the relative error in lesion volumes ϵ(r, *ϑ*) is smallest over all all *ϑ* considered… … … … … … … … … … … … … … .
6. Comparison of final non-dimensional advection-based and perfusionn based temperature fields for *r* = 0.9 mm, *r* = 1.15 mm. The perfusion based temperature was computed using the best estimate perfusion rate *ω*_*b*_. In the advection-based temperature field heating around the vessel downstream of the RF needle decreases with increasing radius.
7. Comparison of final non-dimensional advection-based and perfusion based temperature fields for *r* = 0.9 mm, *r* = 1.15 mm. The perfusion based temperature was computed using the best estimate perfusion rate *ω*_*b*_. In the advection-based temperature field heating around the vessel downstream of the RF needle decreases with increasing radius.
8. Plot of the final objective function 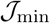, and its two constituent terms versus radius for all the *ϑ* values considered: The colour coding is as follows, Red: *ϑ* = 1e-10, Green: *ϑ* = 1e-9, Blue: *ϑ* = 1e-8… … … … … … … … … … … . .
9. Effect of variation in the perfusion rate *ω* on the perfusion-based lesion volume and relative error for r = 5 mm: The relative error is computed assuming the advection based lesion as the truth… … … … … … … … … … … . .

